# RT-qPCR Validation of Candidate Genes Underlying Thermal Adaptation in *Drosophila subobscura*

**DOI:** 10.1101/2025.07.10.664230

**Authors:** Marta A. Antunes, Margarida Matos, Pedro Simões

## Abstract

Understanding the genetic basis of thermal adaptation is essential in the face of global climate change. In this study, we validate candidate genes implicated in thermal adaptation previously identified through RNA-seq analysis of *Drosophila subobscura* populations experimentally evolved under a progressive warming regime. Using RT-qPCR, we assessed the directionality of gene expression changes—up- or downregulation—across warming and control populations from two latitudinal origins. Out of 27 candidate gene tests, 33.3% showed consistent expression patterns between RNA-seq and RT-qPCR. Our results suggest that effect size, rather than expression level alone, is a key factor driving successful validation. This should be considered when interpreting whole-transcriptome data, which can yield many candidate genes. Future studies should examine how different log2 fold-change thresholds between populations relate to the success rate of validating RNA-seq differential expression results, in order to improve the reliability of candidate gene lists.

## Introduction

Temperature variations significantly influence the physiological and developmental processes of ectothermic organisms, often requiring a robust adaptive response (Urban et al., 2016). The study of thermal adaptation in these organisms is thus relevant in order to provide crucial insights into their responses to that key environmental stressor (Edelsparre et al., 2024; Hoffmann & Sgrò, 2011).

Experimental evolution is an important and increasingly used tool to study adaptation. This approach enables researchers to track changes in phenotypic traits in real time and across multiple generations (Kawecki et al., 2012; Rowiński & Rogell, 2017; Santos et al., 2023). When combined with genome-wide sequencing, it further allows the identification of genetic changes underlying adaptive processes (Santos et al., 2024). Specifically, experimental evolution has proven valuable in studies of thermal adaptation. Among them, several studies have used experimental thermal evolution and RNA-sequencing to investigate the genetic basis of thermal adaptation in Drosophila (see for example the work of Antunes et al., 2024 and Mallard et al., 2018).

While RNA-seq is a powerful tool for discovering differentially expressed genes, complementary validation using other accurate and sensitive techniques is important to confirm these findings. To this end, Reverse Transcription quantitative Polymerase Chain Reaction (RT-qPCR) is a commonly used technique for quantifying gene expression (Chandramohan et al., 2013; Everaert et al., 2017). This method enables accurate measurement of mRNA levels, facilitating the validation of candidate genes identified through high-throughput approaches such as RNA-seq. In a study comparing several RNA-seq pipelines with RT-qPCR, Everaert et al., (2017) found high expression correlations between the two techniques (e.g., qPCR and RNA-seq), but they highlighted systematic discrepancies between quantification technologies in a comparative analysis that ordered genes by rank. In that study, the genes were analyzed all together but also separated by groups of concordant (genes for which the two methods agree on the differential expression status) and non-concordant genes (genes for which the two methods disagree on the differential expression status). Importantly, in that study, differential expression was only considered for genes with log Fold Change >1. Non-concordant genes with ΔFC (difference in log fold change between methods) > 2, as well as those exhibiting opposing directional changes, were predominantly expressed at low levels (Everaert et al., 2017).

*Drosophila subobscura* populations from different latitudes of the species’ European cline evolved in an experimental evolution study under either constant or gradually warming temperatures (Santos et al 2021 Evol). In Antunes et al., (2024), the gene expression response of those populations was analysed. They varied by genetic background, with the high-latitude population showing broader transcriptomic changes. In fact, a total of 4195 candidate genes were identified in high latitude whereas only 949 in low latitude populations. Only 221 genes were differentially expressed in both populations, reinforcing the finding of different adaptive responses between populations from different origins. Many non-essential binding functions were down-regulated in both populations, possibly to conserve energy. Some candidate genes were common among populations. These included genes encoding trichohyalin, larval serum proteins, and fat body proteins, that were all strongly down-regulated in the warming populations. Overall, the results point to both population-specific and shared mechanisms of thermal adaptation (Antunes et al., 2024).

Given what was mentioned before, it would be important to validate the expression of candidate genes detected by Antunes et al., (2024). In the present study, our goal is to validate the expression by RT-qPCR of thermally responsive candidate genes identified in the previously mentioned Antunes et al (2024) study. Our main focus will beoc ng on the direction of expression (i.e. up-regulation or down-regulation comparing experimental populations and their controls). By doing so, we aim to confirm the differential expression patterns and assess the reliability of the selected candidate genes for thermal adaptation. We defined several criteria to select the genes to be tested, namely those with strong evidence for thermal selection (through high significance levels or log2FC values) or consistency between latitudinal populations (see details below).

## Materials and Methods

### Population maintenance and thermal selection regimes

In 2013, a total of 234 and 160 female flies were collected from Adraga (PT, Portugal) and Groningen (NL, The Netherlands), respectively. By the fourth generation in the laboratory, the populations were three-fold replicated (PT1-3 and NL1-3) and maintained in discrete generations under a synchronized 28-day cycle, a 12L:12D photoperiod, and a constant temperature of 18ºC, with egg collection for the next generation around 8 days of the imago’s age (Simões et al., 2017).

In 2019, a thermal selection regime was implemented in populations of both origins: the global warming regime (WNL and WPT) (Santos et al., 2021). This regime includes daily temperature fluctuations, ranging between 15ºC and 21ºC in the first generation, with a progressive increase in both daily mean temperature (0.18 °C) and amplitude (0.54 °C) per generation until generation 22 (see Santos et al., 2023). Simultaneously, the ancestral PT and NL populations were maintained as controls (C) at a constant temperature of 18ºC (the control temperature).

### Samples for RNA-seq analysis and candidate genes definition

Our focus is to try to validate several candidate genes obtained in Antunes et al. (2024). As there described, after 23 generations of evolution under the warming selection regime, flies from that regime were placed for one generation in a common garden environment at the control temperature. Eggs from the derived flies as well as from the control regime were then exposed to the temperature of the warming regime where they developed, allowing for sample collection. The study analyzed 12 pooled samples representing two selective regimes (control vs. warming), two latitudinal origins (from here on called histories: high vs. low), and three replicate populations per regime^*^history (e.g., NL1-3). Each sample consisted of a pool of 45 adult females, sex-screened two days before the flash-frozen, that took place at their 8th day of adulthood. Total RNA was extracted from each sample, the highest-quality RNA extracts were used for sequencing while remaining RNA elutions were stored for further analysis. Candidate genes were identified by comparing gene expression of the selective regimes with the respective controls where genes with a p-value < 0.001 were considered candidate genes. For RT-qPCR validation, the second-best RNA extract from the same pooled samples was used (see Supplementary Table S1), allowing to test the reproducibility of differential expression patterns and confirm candidate gene responses (see details below).

### Gene validation and criteria

The reference gene used in this work is the elongation factor 1-alpha 2 (eEF1alpha2), which has good expression stability associated with temperature in *Drosophila melanogaster* (Lü et al., 2018). A set of candidate genes implicated in thermal adaptation from Antunes et al. (2024) were selected to be validated by RT-qPCR. The criteria were the following: 1) Candidate genes under selection across low and high latitude populations in both control and warming environments - 9 genes; and 2) 2 Genes with highly significant p-values for selection in warming environment and 2 gens with highest |log2FC| - 4 genes. 3) Genes previously linked to thermal selection in other thermal E&R studies (e.g.Santos et al., 2024) - 14 genes. The complete list of candidate genes analysed in this study is provided in Supplementary Table S2.

### Complementary DNA (cDNA) synthesis

cDNA synthesis was done according to Xpert cDNA Synthesis kit protocol (GRISP). Specifically, 4ul of 5x Reaction buffer,1 ul of dNTP mix, 1ul of hexaprimer and 1ul of Xpert RTase per sample were added to a RNase-free microtube kept on ice. 7ul of the mix were distributed in each RNase-free microtube, as well as the volume corresponding to 50ng of template RNA and the remaining volume of RNase free water. The final volume in each microtube was 18ul.

### Primer design and Real time quantitative polymerase chain reaction (RT-qPCR)

Primer design was performed using Primer-BLAST with the following parameters: a maximum PCR product size of 200 bp, and, when possible, the option to ensure primers spanning an exon-exon junction. Primer melting temperatures were set to a minimum of 57°C, an optimum of 60°C, and a maximum of 63°C (see the primers that were used in this work, Supplementary Table S3).

The RT-qPCR mixes were prepared according to Supplementary Table S3. The cDNA was diluted to a final concentration of 20 ng. The RT-qPCR reaction plates were run on the Bio-Rad CFX96 qPCR system, using the qPCR cycling according to manufacturer instructions (GRISP, details in Supplementary Table S4).

### Comparison between RNA-seq and RT-qPCR data

We used the following formula to compare the results of RT-qPCR with those of RNA-seq: - ΔΔCq=(Cqreference gene-Cqtarget gene) warming samples-(Cqreference gene-Cqtarget gene) control samples (see Applied Biosystems guide, 2008). In this formula Cq is the value obtained after the thermocycler run and corresponds to the number of cycles at which the fluorescence log reaches the threshold level. The metrics used for RNA-seq was the log2Fold Change in which the fold change is the ratio between the normalized counts in the warming (evolved) sample and the normalized counts in the control sample (Chen et al., 2018). We considered validated genes when the sign of both metrics match in all or at least two of the three replicate populations of the corresponding history. Also, in a less stringent analysis we considered concordant genes when the sign of the average of the three replicates matches between methods (details below).

## Results

### Overall validation of gene expression

In Figure 1 we summarize the results of the validation of the genes analyzed by RT-qPCR. The green segment represents genes that exhibited the same signal in RT-qPCR and RNA-seq (i.e. up or down regulated, comparing warming and control populations) across all replicates, while the yellow section includes genes that in RT-qPCR showed two correct signals (that is, the same as in RNA-seq) but one weakly incorrect signal (that is very low signal of opposite direction of regulation), indicating partial validation. Considering the two categories together, in low latitude populations 3 out of 10 genes (30%) were validated while in high latitude populations there were 6 validated genes out of 17 (35.3%) – see Figure 1. Thus, pooling the results of the populations of both histories together we had a successful validation for 9 out of 27 tests (33.3%), as illustrated in the rightmost pie chart of Figure 1. Validation of expression of each specific candidate gene is provided in Supplementary Table S2. The last column of that table indicates which genes were validated, either fully or partially, corresponding to those marked respectively in green or yellow in Figure 1. Validation by different methods, namely using as reference the mean of total genes in each sample, did not show an improvement in the validation percentages (Supplementary Figure S1).

**Figure 1.**
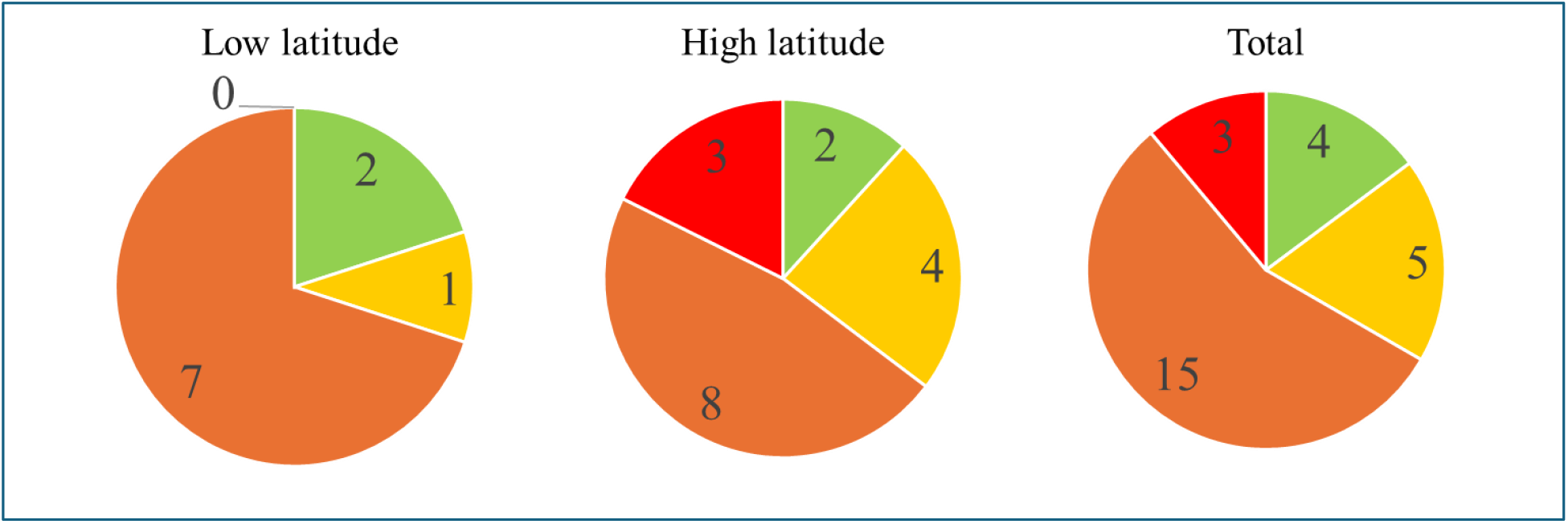
Pie charts showing the number of tests categorized by signal of expression accuracy. Green represents tests with a correct signal in all replicates, yellow indicates tests with two correct signals and one weak incorrect signal, orange represents tests with either two incorrect signals or one very strong incorrect signal and red shows tests with three incorrect signals. The left pie chart refers to low latitude populations, in the middle is the pie chart for high latitude populations and on the right the pie chart with the total of observations (genes in both latitudinal populations).

### Validation by criteria

Looking at the criteria of selection of candidate genes, we had a successful validation of 33 % of tested genes (considering independent tests cases where the same genes were tested in both histories) in criteria 1, 50% in criteria 2 and 29% in criteria 3, to an overall, total of 33% of tests validating the corresponding genes (see details in Table 1 and discriminated by history in Supplementary Table S5). The higher percentage of validation for the second criteria (when p-value is lower or Log2FC is higher) raised whether it is associated with a higher log2FC between thermal regimes. To clarify this, we focused on the patterns observed for all genes showing high Log2FC (>2). Two validated genes recruited by other criteria were considered, as they also showed high Log2FC: larval serum protein is included in criteria 1 and was validated whereas trichohyalin is included in criteria 2 and was also validated. When taking into account the Log2FC levels, we observed 22% of validated genes have a log2FC higher than 2 whereas 78% have a log2FC lower than 2, and 6% of non-validated genes have log2FC higher than 2 whereas 94% have a log2FC lower than 2. Overall, there is a higher validation percentage when the log2FC is higher than 2 (from 6% to 22%). Considering now only candidate genes with log2Fc higher than 2, 67% were validated whereas only 30% were validated when log2FC is lower than 2. Nevertheless, it is important to take into account the small number of genes with such high log2FC (n=3). Supplementary Table S5 presents the validation analysis stratified by criteria but also by history of the populations. A higher validation percentage is observed for candidate genes in high-latitude populations under criterion 1 (50 vs 20), although the number of genes analyzed in this category is low. For the remaining criteria, validation percentages are relatively similar.

**Table 1.** Number of tests categorized by signal of expression accuracy, discriminated by criteria. Criteria 1 corresponds to candidate genes across low and high latitude populations in both control and warming environments, criteria 2 includes genes with highly significant p-values for selection in warming environment or highest |log2FC and criteria 3 corresponds to candidate genes that were also linked to thermal selection in other thermal E&R studies (e.g. Santos et al., 2024) See explanation of colours in figure 1.

|  | nr of genes | nr of analyses | A | B | C | D | val % (A+B) |
| --- | --- | --- | --- | --- | --- | --- | --- |
| **criteria 1** | 5 | 9 | 3 | 0 | 5 | 1 | 33 |
| **criteria 2** | 3 | 4 | 1 | 1 | 2 | 0 | 50 |
| **criteria 3** | 12 | 14 | 0 | 4 | 8 | 2 | 29 |
| **total** | 20 | 27 | 4 | 5 | 15 | 3 | 33 |

### Analysis of Concordance of gene expression

We also assessed the concordance in the direction of gene expression changes between RNA-seq and RT-qPCR results for candidate gene expression across populations from different latitudinal regions (Figure 2). In this case we only looked at the average value of the three replicate populations for both RT-PCR and RNA seq data, not accounting for differences between replicate populations. With this less stringent analysis we found moderate concordance between the two techniques, with 60% of genes in low-latitude populations and 65% in high-latitude populations displaying upregulation or downregulation across both data sets (i.e., falling in the first or third quadrants, see Figure 2).

**Figure 2.**
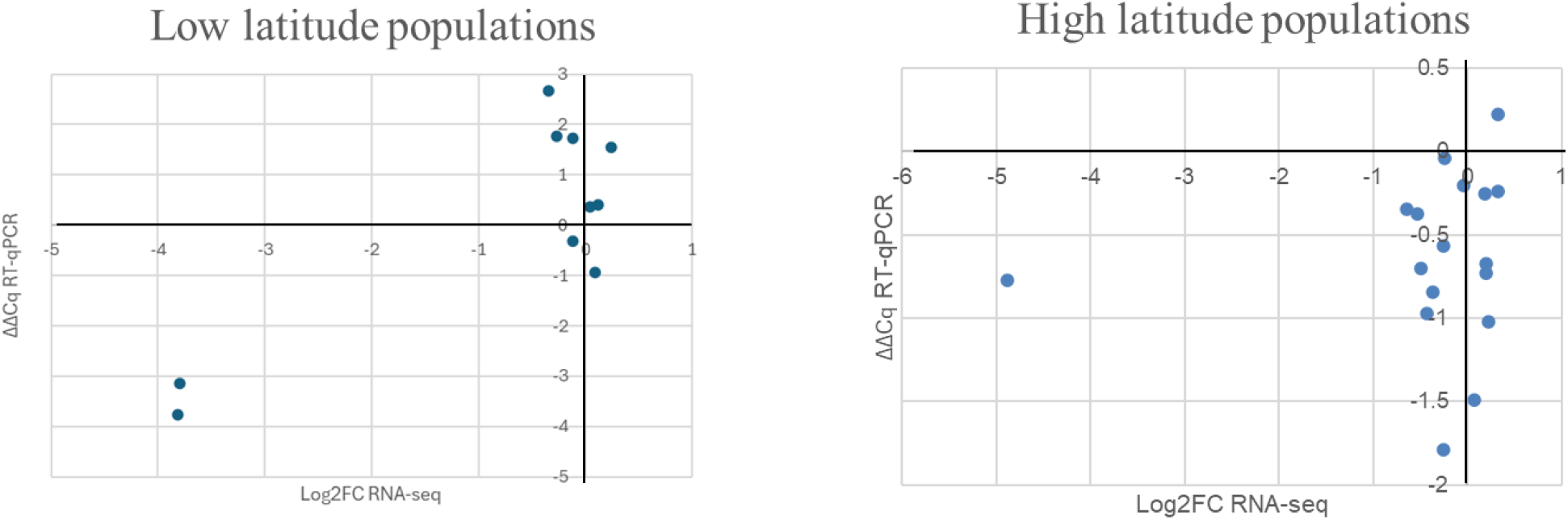
Plots showing concordance or discordance in expression between RT-qPCR and RNA-seq methods. Each point represents a candidate gene. The x-axis shows the log2 fold change (Log2FC) in gene expression based on RNA sequencing (RNA-seq), while the y-axis represents the ΔΔCq values obtained from RT-qPCR. Genes that fall in the first or third quadrants indicate the same sign of expression - concordance - between RNA-seq and RT-qPCR measurements, meaning both methods show upregulation or downregulation of the gene. Genes in the second or fourth quadrants show discordance between the two methods.

## Discussion

The study of gene expression allows for a deeper understanding of the process of adaptation to global warming (Oomen & Hutchings, 2022). Here we provide a report on the validation of some candidate genes for thermal adaptation, the latter obtained from the RNA-seq study of the complete transcriptome of experimentally-evolved *Drosophila subobscura* populations (Antunes et al., 2024). We were able to validate the expression of some candidate genes (around 33.3% of tests across latitudinal populations) and observed a moderate concordance in the direction of expression changes (up vs downregulation) when comparing both techniques but also found discrepancies.

Several factors may have contributed to this generally low validation. First, RNA quality can influence transcript quantification (Vermeulen et al., 2011). In our study, the RNA used for RT-qPCR validation was of poorer quality compared to the RNA used for sequencing. Another possibility would be that low validation can be associated with low general expression of the candidate genes (Asmann et al., 2009; Everaert et al., 2017). It might be expected that higher gene expression would contribute to higher Log2FC. In fact, the criteria with higher validation was criteria 2 (reflecting high Log2FC or low p-values). In fact, our results do suggest that there is a higher probability of validation when the magnitude of change between thermal regimes is high, with 2 out of 3 genes with log2FC higher than 2 being validated. Nevertheless, this increased validation is not due to higher gene expression levels of such genes, as might be expected from the findings of Everaert et al 2017. Such genes had expression levels that were below the mean expression values across candidate genes.

It is likely that the low levels of general validation in our study might have resulted from the rather low number of genes with high magnitude of effect size between thermal selection regimes (i.e. high log2FC). While it could be argued that only genes with high log2FC should have been retained, we chose to include all significantly differentially expressed genes—regardless of the magnitude of change— to capture not only large expression shifts but also more subtle changes, though consistent across replicate populations, that may provide a broader view of thermal adaptation to impact of global warming. In any case we need to be cautious as to generalizations, given the low number of genes with high log2FC analysed in this work. It has been shown that the quality of the reads in RNA-seq can also have an influence on the quantification. Specifically, low quality reads (below Q20) are more frequent in genes that show discrepancies between techniques (Everaert et al. 2017). In the case of our study, these factors are unlikely to have affected the analysis, as we conducted quality control, where only reads with a quality score above Q20 were retained.

The choice of reference gene is also important. Different results have been reported depending on the reference gene or genes used in the validation (Lü et al., 2018; Shakeel et al., 2018; Shu et al., 2018). Given the above, it is of course possible that validation percentages obtained with other reference gene or genes could differ from those reported here. Additionally, alternative normalization approaches—such as using the mean expression of all genes in each sample (Everaert et al., 2017; Vinje & Friedman, 2023) or the mean expression of non-candidate genes—were also considered. However, in this study, these methods did not improve validation performance and proved suboptimal.

Despite these challenges, genes such as larval serum protein and trichohyalin were successfully validated by RT-qPCR. These genes were highlighted as candidate genes for thermal adaptation in Antunes et al. (2024) because of their consistent down-regulation in both warming populations, of low and high latitudinal origin. These results strengthen the reliability of at least some of the genes highlighted by us in our transcriptomic analysis.

To summarize, our study suggests that the effect size of expression changes between populations is likely to be a key aspect in determining the efficacy of validation of RNAseq results. This should be taken into consideration when interpreting such whole-transcriptome data, that potentially generates a very high number of candidate transcripts. Moving forward, a more detailed investigation into log2FC thresholds between populations and their association with the efficiency of validation of RNA-seq results of differential expression would be helpful to increase reliability of candidate gene lists derived from gene expression studies.

## Notes

### Competing Interest Statement

The authors have declared no competing interest.

### Funding

- Fundação para a Ciência e Tecnologia: 2020.09172.BD, PTDC/BIA-EVL/28298/2017, UID/00329/2025

