## Supplementary tables and figures for "RT-qPCR Validation of Candidate Genes Underlying Thermal Adaptation in *Drosophila subobscura*": Supplementary Tables-CAP5.docx

Table S1 - Quantification and ratios of RNA samples used for RT-qPCR validation obtained by Nanodrop.

| sample | C(ng/μl) | 260/280 ratio | 260/230 ratio |
| --- | --- | --- | --- |
| PT1 | 199.0 | 2.16 | 2.40 |
| NL1 | 22.8 | 2.23 | 1.50 |
| WPT1 | 256.4 | 2.14 | 2.17 |
| WNL1 | 57.6 | 2.17 | 1.83 |
| PT2 | 230.9 | 2.14 | 2.36 |
| NL2 | 65.9 | 2.16 | 2.26 |
| WPT2 | 384.7 | 2.13 | 1.38 |
| WNL2 | 23.6 | 2.07 | 1.56 |
| PT3 | 64.0 | 2.12 | 1.87 |
| NL3 | 139.2 | 2.17 | 2.20 |
| WPT3 | 271.8 | 2.13 | 2.38 |
| WNL3 | 123.2 | 2.03 | 2.47 |

Table S2 – List of genes and respective primers for RT-qPCR.

| Criteria | gene ID | Population in which is candidate | annotation | Forward primer | Reverse primer | Validation status |
| --- | --- | --- | --- | --- | --- | --- |
|  | LOC117899650 (ref) |  | elongation factor 1-alpha 2 | CCTCCTGAAGCCAGGAATGG | CCACGGGGTGGATTGTTCTT |  |
| Gene under selection across low and high latitude populations in both control and warming environments | LOC117902204 (both pops) | Both | larval serum protein 1 gamma chain | GACAAGGCCCAGTACAAGGAAT | GGGTCTTCACCAGAGAACCG | Validated for low |
|  | LOC117898973 (both pops) | Both | probable ATP-dependent RNA helicase spindle-E | GCGCATGTACTTGACCTACCT | TAGGTAATCGGCACCAGGGA | Validated for high |
|  | LOC117901038 (both pops) | Both | fibroblast growth factor receptor substrate 2 | GATCAACCCATCCCGGAACC | GAAAGACGTTGTCGTGACGC | Not validated |
|  | LOC117891209 (both pops) | both | LOW QUALITY PROTEIN: protein lethal(2)denticleless | AGAGCATGGCTTCTCGAACG | TCGTTGGCAATGGCCAGTAT | Not validated |
|  | LOC117898405 (both pops) | Both | transforming growth factor-beta-induced protein ig-h3, temos varias isoformas | CGTGAGTGAGTGACACCCC | CTAGTGGTATGGGCGCGAAC | Validated for high |
| very low p-value | LOC117894143 (both pops) | Both | heat shock protein 27(verificar isoforma) | AAGAATCGCGAGTGGACGTT | TTGTTCGTAGCCGGAACCAG | Validated for high |
| high log2FC | LOC117892218 (both pops) | Both | trichohyalin | ACTTGCATTGCCCGTAGGTT | GGAGGTGTGTGGGACATGAG | Validated for low |
| Gene previously linked to thermal selection in other thermal E&R studies (e.g. Santos et al. 2024 MBE) and also defined as candidates in our study | LOC117896292 (high lat pops) | high lat | pyruvate dehydrogenase E1 component subunit beta, mitochondrial | GACGGTGCCTACAAGGTATCC | AGGTCATGAACTCGCAGACG | Not validated |
|  | LOC117902220 (high lat pops) |  | DNA-directed RNA polymerases I and III subunit RPAC1 | TTCAAGTGGCCATCATTCGAAAC | ACCTCACTGAGCATCAAGCG | Not validated |
|  | LOC117889582 (low lat pops) |  | probable ATP-dependent RNA helicase DHX34 | GACCAGCAATATCCCTCCACC | GTCTTGCTCGGTTAGTGCCA | validated |
|  | LOC117890021 (both pops) |  | ornithine decarboxylase antizyme | TTCCGCACCATCTCGACTTC | TTACGGCTGTAGCCCGTTAAG | Not validated |
|  | LOC117903461 (high lat pops) |  | vinculin | CTGTCGCCCAGCAGGTATC | CTGTTGATGGTCTCCCGTCC | Not validated |
|  | LOC117889583 (high lat pops) |  | N-alpha-acetyltransferase 16, NatA auxiliary subunit | GTGAGCAGAGACAAAGCGGA | GCATGGGCGAGTTTCCCCTA | Not validated |
|  | LOC117890527 (high lat pops) |  | Drosophila subobscura T-complex protein 1 subunit epsilon | TTCCTGGTACGTTCGCGTTT | TGCCGCCATAATATGCGTCTTT | Not validated |
|  | LOC117890693 (high lat pops) |  | hsc70-interacting protein 1-like | ACTCCGAGGGAGTCAGCATC | CTGGCTGTATAACGCCTTCCA | Validated |
|  | LOC117895027 (high lat pops) |  | L-lactate dehydrogenase | AACAAGTGGTGGACTCTGCC | CGATGCCATGTTCACCCAGT | Not validated |
|  | LOC117903707 (both pops) | Both | pyruvate dehydrogenase E1 component subunit alpha | TGCAAGTGAATCGCCCCTTC | CGCCTCATCCTTGGTCAGTT | Validated for high |
|  | LOC117896570 (high lat pops) |  | 5'-AMP-activated protein kinase catalytic subunit alpha-2 | CGTCAAGGTCGCTGTCAAGA | GAAGGAGTGGAGATCACCTGGTA | Not validated |
|  | LOC117898834 (high lat pops) |  | 5'-AMP-activated protein kinase subunit gamma-1 | CCCTTGGAGAGCATCAAAGAGT | ATCTGTGAGTCGTCCTCTTCTTTC | Validated |

Table S3 – Composition of the master mixes for RT-qPCR.

| Component | 1x reaction |
| --- | --- |
| Xpert Fast SYBR 2x master mix Blue | 10μl |
| Forward primer | 1μl |
| Reverse primer | 1μl |
| Water | 2μl |
| cDNA | 6μl |
| Vfinal (RT-qPCR reation) | 20μl |

Table S4 – Table with qPCR temperature cycles

| Number of cycles | Temperature | Time |
| --- | --- | --- |
| 1x | 95ºC | 2min |
| 40x | 95ºC | 5sec |
|  | 60ºC | 20sec |

Table S5 -Table with validation per criteria and origin of populations.

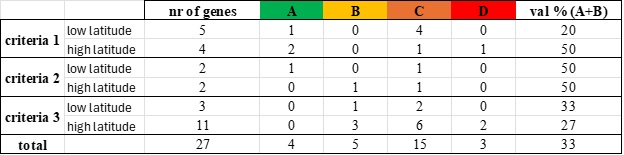

**Supplementary Figures**

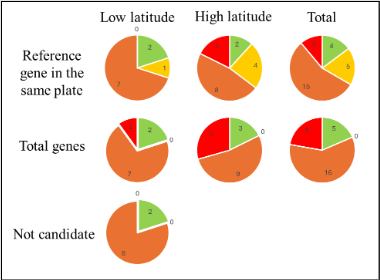

Figure S1 - Pie charts with validation using different methods. In the 3 upper pie charts, results of the method with a reference gene are presented, in the 3 middle pie charts are the results using as reference the mean of total genes analysed by RT-PCR method in each sample, and finally, in the bottom pie chart the results using as reference the non-candidate genes in the population under study (e.g. non-candidate genes in low latitude populations but candidates in high letitude populations).
